# The Gene-Rich Genome of the Scallop *Pecten maximus*

**DOI:** 10.1101/2020.01.08.887828

**Authors:** Nathan J Kenny, Shane A McCarthy, Olga Dudchenko, Katherine James, Emma Betteridge, Craig Corton, Jale Dolucan, Dan Mead, Karen Oliver, Arina D Omer, Sarah Pelan, Yan Ryan, Ying Sims, Jason Skelton, Michelle Smith, James Torrance, David Weisz, Anil Wipat, Erez L Aiden, Kerstin Howe, Suzanne T Williams

## Abstract

**Background:** The King Scallop, *Pecten maximus*, is distributed in shallow waters along the Atlantic coast of Europe. It forms the basis of a valuable commercial fishery and its ubiquity means that it plays a key role in coastal ecosystems and food webs. Like other filter feeding bivalves it can accumulate potent phytotoxins, to which it has evolved some immunity. The molecular origins of this immunity are of interest to evolutionary biologists, pharmaceutical companies and fisheries management.

**Findings:** Here we report the genome sequencing of this species, conducted as part of the Wellcome Sanger 25 Genomes Project. This genome was assembled from PacBio reads and scaffolded with 10x Chromium and Hi-C data, and its 3,983 scaffolds have an N50 of 44.8 Mb (longest scaffold 60.1 Mb), with 92% of the assembly sequence contained in 19 scaffolds, corresponding to the 19 chromosomes found in this species. The total assembly spans 918.3 Mb, and is the best-scaffolded marine bivalve genome published to date, exhibiting 95.5% recovery of the metazoan BUSCO set. Gene annotation resulted in 67,741 gene models. Analysis of gene content revealed large numbers of gene duplicates, as previously seen in bivalves, with little gene loss, in comparison with the sequenced genomes of other marine bivalve species.

**Conclusions:** The genome assembly of *Pecten maximus* and its annotated gene set provide a high-quality platform for a wide range of investigations, including studies on such disparate topics as shell biomineralization, pigmentation, vision and resistance to algal toxins. As a result of our findings we highlight the sodium channel gene *Nav1*, known as a gene conferring resistance to saxitoxin and tetrodotoxin, as a candidate for further studies investigating immunity to domoic acid.

## Data Description

### Context

Scallops are bivalve molluscs (Pectinida, Pectinoidea, Pectinidae; Fig. 1A, B), found globally in shallow marine waters, where their filter-feeding lifestyle helps perform a variety of ecological functions [1]. There are around 400 living scallop species [2], and of these, *Pecten maximus* (Fig. 1A), also known as the King Scallop, Great Scallop and St James Scallop, is perhaps the best-studied European species. *Pecten maximus* is found around the coast of western Europe from northern Norway to the Iberian Peninsula (Fig. 1C) where it is locally common in many areas, and it can occasionally be found more distantly in West Africa and on mid-North Atlantic islands [2]. It is commercially fished across its range, most heavily around France and the United Kingdom [3, 4] and is the most valuable single species fishery in the English Channel with around 35,000 tonnes of international landings reported in 2016 [4]. It has also been cultivated in aquaculture, particularly in the United Kingdom, Spain, Norway and France, although with limited commercial production [5, 6]. It is an important part of the ecosystems within which it occurs, performing key roles in food webs, both as a prey species and more indirectly by cycling nutrients when filter feeding [1].

**Figure 1:**
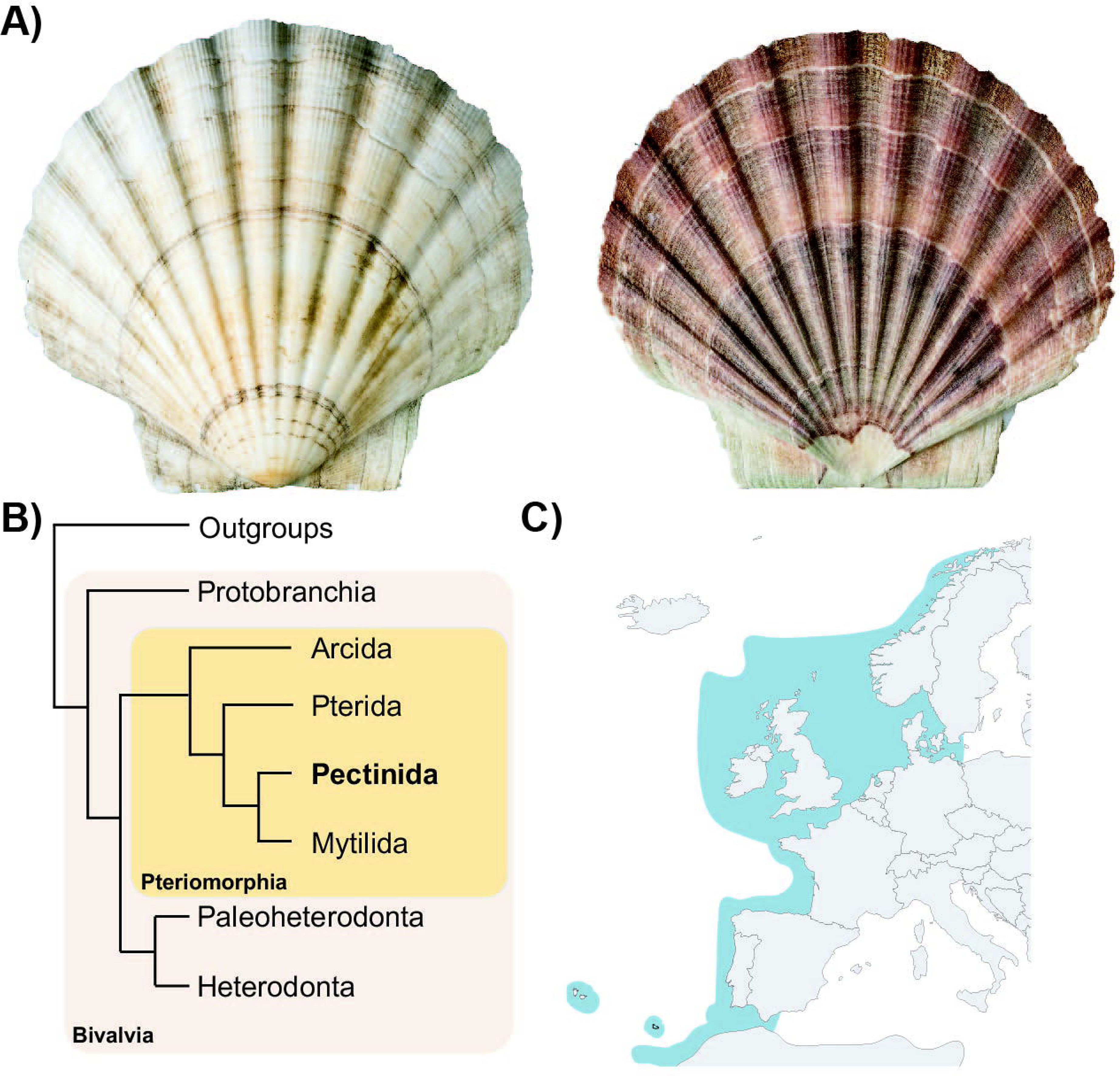
A) Photo of both valves of the shell of *Pecten maximus*, from the specimen sequenced in this work (NHMUK 20170376). B) Diagrammatic cladogram illustrating the phylogeny of the Bivalvia [after 65], showing the major sub-classes of Bivalvia and (internally boxed) the major divisions of the Pteriomorphia. *Pecten maximus* is a member of the superfamily Pectinoidea which includes Pectinidae (scallops), Propeamussiidae (glass scallops) and Spondylidae (spiny oysters) and together with their close relatives (Anomioidea, jingle shells; Dimyoidea, dimyarian oysters; and Plicatuloidea, kittenpaw clams) these superfamilies form the Order Pectinida. C) Distribution map of *P. maximus*, showing range (dark blue) of species across Northern Europe and surroundings [Map from simplemaps, distribution according to 2].

Previous studies in this species have aimed to understand its population dynamics, swimming behaviour, visual systems and reproduction. Of particular interest to medicine, fisheries management and molecular biology is the means by which this species is immune to neurotoxins like saxitoxin (STX) and domoic acid (DA). DA and STX are potent neurotoxins produced by certain species of phytoplankton, including dinoflagellates and diatoms, which may be present in large blooms [3]. Some shellfish (e.g. scallops, *P. maximus;* mussels, *Mytilus edulis*; cockles, *Cerastoderma edule*; razor clams, *Siliqua patula*), fish (e.g. anchovy, *Engraulis mordax*; European sardine, *Sardina pilchardus*; and Pacific Halibut, *Hippoglossus stenolepis*), and crabs (e.g. *Cancer magister*) accumulate algal neurotoxins by filtration of phytoplankton or by ingestion of contaminated organisms, with species-specific accumulation rates [7–9]. In humans, ingestion of DA or STX has been associated with gastrointestinal and neurological symptoms [10, 11]. In severe cases, poisoning by DA may lead to death or permanent memory loss, a syndrome known as Amnesic Shellfish Poisoning and in the case of STX, paralysis (Paralytic Shellfish Poisoning) [12]. Curiously, however, shellfish and fish that routinely accumulate algal toxins are often able to do so without apparent effect on their health [13, 14]. The resistance of *P. maximus* in particular, and of bivalve molluscs more generally, to these potent toxins is of keen interest to fisheries groups, health care providers and molecular biologists, yet the genetic mechanism behind this remains unknown. Detailed investigation into this phenomenon, along with many others, would be greatly aided by a genome resource.

At the time of writing, nine bivalve genomes are available, with those of the Pacific oyster *Crassostrea gigas* [15] and the pearl oyster *Pinctada fucata* [16] in particular having been used for a variety of investigations into bivalve biology. Scallops have been the subject of genome sequencing projects in the past, with genomes published for three species, *Azumapecten farreri* (as *Chlamys* [17]) and *Mizuhopecten yessoensis* (as *Patinopecten*; [18]) from the subfamily Pedinae, and *Argopecten purpuratus* from the subfamily Pectininae [19]. Other sequenced genomes for more distantly related bivalves include those of the Sydney Rock Oyster *Saccostrea glomerata* [20], Eastern oyster *Crassostrea virginica* [unpublished, but see 21], the Snout Otter Clam *Lutraria rhynchaena* [22], Blood Clam *Scapharca broughtonii* [23] and Manila Clam *Ruditapes philippinarum* [24]. There are also extant resources for the mussel *Mytillus galloprovincialis* [25] and the freshwater mussels *Venustaconcha ellipsiformis* [26], *Limnoperna fortunei* [27], *Dreissena rostriformis* [28] and *Dreissena polymorpha* [unpublished, but see 29]. Of these resources, only the assemblies for *Crassostrea virginica* and *Scapharca broughtonii* are of chromosomal quality, and the scaffold N50 of the other resources varies widely.

These studies demonstrate that bivalve genomes are often 1 Gbp or more in size, and generally exhibit large amounts of heterozygosity, related to their tendency to be broadcast spawners with excellent dispersal capabilities, resulting in large degrees of panmixia. Gene expansion has been noted as a characteristic of the clade, with some species exhibiting tandem duplications and gene family expansions, particularly in genes associated with shell formation and physiology (e.g. HSP70 [30]).

Here we describe the genome of the King Scallop, *Pecten maximus*, which has been assembled from PacBio, 10x Genomics and Hi-C libraries. It is a well-assembled and complete resource, and possesses a particularly large gene set, with duplicated genes comprising a substantial part of this complement. This genome and gene set will be useful for a range of investigations in evolutionary genomics, aquaculture, population genetics, and the evolution of novelties such as eyes and colouration, for many years to come.

### Methods

#### Sample information, DNA extraction, Library Construction, Sequencing and Quality Control

A single adult *Pecten maximus* was purchased commercially, marked as having been collected in Scotland. The shell was preserved and is deposited in the Natural History Museum, London with voucher number NHMUK 20170376. The adductor muscle was used for high molecular weight DNA extraction using a modified agarose plug based extraction protocol. DNA was cleaned using a standard phenol/chloroform protocol, concentration determined with a Qubit high sensitivity kit, and high molecular weight content confirmed by running on a Femto Pulse (Agilent, Santa Clara, USA).

PacBio and 10x Genomics linked read libraries were made at the Wellcome Sanger Institute High-Throughput DNA Sequencing Centre by the Sanger Institute R&D and pipeline teams using established protocols. PacBio libraries were made using the SMRTbell Template Prep Kit 1.0 and 10x libraries using the Chromium Genome Reagent Kit (v2 Chemistry). These libraries were then sequenced on Sequel 1 and Illumina HiSeq X Ten platforms respectively at the Wellcome Sanger Institute High-Throughput DNA Sequencing Centre. The raw data are available from ENA at: https://www.ebi.ac.uk/ena/data/view/ERS3230380. Hi-C reads were created by the DNA Zoo Consortium (www.dnazoo.org), and submitted to NCBI with accession number SRX6848914. Read quality, adapter trimming and read length was assayed using NanoPlot and NanoComp (PacBio reads) [31] and FastQC (10x) (Supplementary File 1). PacBio libraries provided approximately 65.9x coverage of this genome, 10x reads and Hi-C provided a further 113.7x and 63.4x estimated coverage, respectively, assuming a genome size of 1.15 Gbp. A summary of statistics relating to these reads can be found in Table 1.

**Table 1:** Libraries sequenced and used in assembly, with accession numbers as shown.

| **Library Type** | **Number of sequencing runs** | **Number of reads** | **Number of bases (Gbp)** | **GC%** | **Nominal Coverage (1.15 Gbp genome)** | **Accessions** |
| --- | --- | --- | --- | --- | --- | --- |
| 10x | 4 | 433,117,392 | 130.8 | 39.5 | 113.7x | ERR3316025-ERR3316028 |
| PacBio | 13 | 7,246,290 | 75.8 | 38.99 | 65.9x | ERR3130278-ERR3130281,<br>ERR3130284-ERR3130292 |
| Hi-C | 1 | 241,297,364 | 72.9 | 38.7 | 63.4x | SRX6848914 |

#### Genome Assembly

PacBio reads were first assembled with wtdbg2 v2.2 using the `-xsq` preset option for PacBio Sequel data [32]. The PacBio reads were then used to polish the contigs using Arrow (genomicconsensus package, PacBio tools). This was followed by a round of Illumina polishing using the 10X data which consisted of aligning the 10X data to the contigs with longranger align, calling variants with Freebayes 1.3.1 [33] and applying homozygous non-reference edits to the assembly using bcftools-consensus (https://github.com/VGP/vgp-assembly/tree/master/pipeline/freebayes-polish). Scaffolding was performed using scaff10x 4.2 (https://github.com/wtsi-hpag/Scaff10X). Hi-C based scaffolding was performed by the DNA Zoo Consortium using 3D-DNA [34], followed by manual curation using Juicebox Assembly Tools [35]. A further round of polishing with Arrow was performed on the resulting scaffolds, with reads spanning gaps contributing to filling in assembly gaps. This was followed by a further two rounds of FreeBayes Illumina polishing. Finally, the assembly was analysed and manually improved using gEVAL [36].

Full statistics regarding our assembly can be seen in Table 2. The assembly contains a total of 918,306,378 bp, across 3,983 scaffolds. The N50 is 44,824,366 bp, with 50% of the genome found in 10 scaffolds. The Hi-C analysis identified *P. maximus* possesses 19 pairs of chromosomes, in agreement with prior studies [37], and these are well recovered in our assembly, with 844,299,368 bp (92%) of our assembly in the 19 biggest scaffolds, the smallest of which is 32,483,354 bp, and the largest is 60,076,705 bp in length; only 0.08% of the assembly are represented as Ns (691,874bp). The assembly was screened for trailing Ns, and for contamination against databases of common contamination sources, adaptor sequences and organelle genomes derived from NCBI (using megaBLAST algorithm, requiring e-value <= 1e-4, sequence identity >= 90%, and for genome comparisons, match length >= 500). This process identified no contamination. The Hi-C contact map for the final assembly is shown in Fig 2D, and demonstrates the integrity of the chromosomal units. The interactive version of the contact map is available at http://bit.ly/2QaYqvk (powered by Juicebox.js [38]) and on the http://www.dnazoo.org/assemblies/Pecten_maximus webpage. Our assembly is the most contiguous of all published bivalve genome assemblies to date (Table 3).

**Table 2:** Basic metrics relating to assembled genome.

|  |  |
| --- | --- |
| **Total assembly length [bp]** | 918,306,378 |
| **GC Content of scaffolds** | 36.62% |
| **Max scaffold length [bp]** | 60,076,705 |
| **N50 scaffold length [bp]** | 44,824,366 |
| **N90 scaffold length [bp]** | 32,483,354 |
| **Number of scaffolds** | 3,983 |
| **Number of scaffolds in N50** | 10 |
| **Number of chromosomes** | 19 |
| **% genome, chromosome-length scaffolds** | 92% |
| **N content, total [bp]** | 691,874 |

**Table 3.**
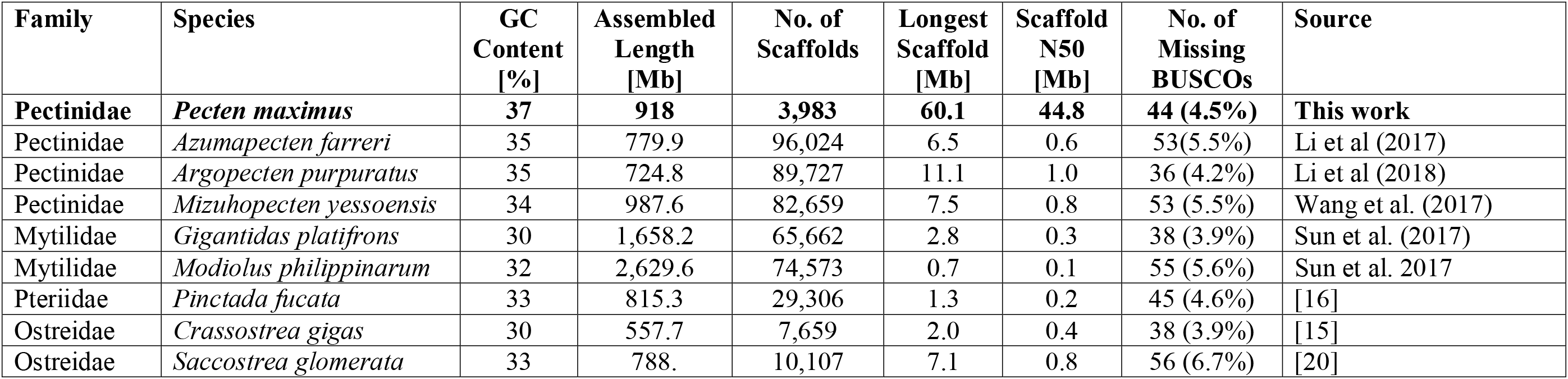
Genomic assemblies of a number of marine bivalves, and summary statistics relating to these assemblies (data in [71]).

| Family | Species | GC Content [%] | Assembled Length [Mb] | No. of Scaffolds | Longest Scaffold [Mb] | Scaffold N50 [Mb] | No. of Missing BUSCOs | Source |
| --- | --- | --- | --- | --- | --- | --- | --- | --- |
| **Pectinidae** | ***Pecten maximus*** | **37** | **918** | **3,983** | **60.1** | **44.8** | **44 (4.5%)** | **This work** |
| Pectinidae | *Azumapecten farreri* | 35 | 779.9 | 96,024 | 6.5 | 0.6 | 53(5.5%) | Li et al (2017) |
| Pectinidae | *Argopecten purpuratus* | 35 | 724.8 | 89,727 | 11.1 | 1.0 | 36 (4.2%) | Li et al (2018) |
| Pectinidae | *Mizuhopecten yessoensis* | 34 | 987.6 | 82,659 | 7.5 | 0.8 | 53 (5.5%) | Wang et al. (2017) |
| Mytilidae | *Gigantidas platifrons* | 30 | 1,658.2 | 65,662 | 2.8 | 0.3 | 38 (3.9%) | Sun et al. (2017) |
| Mytilidae | *Modiolus philippinarum* | 32 | 2,629.6 | 74,573 | 0.7 | 0.1 | 55 (5.6%) | Sun et al. 2017 |
| Pteriidae | *Pinctada fucata* | 33 | 815.3 | 29,306 | 1.3 | 0.2 | 45 (4.6%) | [16] |
| Ostreidae | *Crassostrea gigas* | 30 | 557.7 | 7,659 | 2.0 | 0.4 | 38 (3.9%) | [15] |
| Ostreidae | *Saccostrea glomerata* | 33 | 788. | 10,107 | 7.1 | 0.8 | 56 (6.7%) | [20] |

**Figure 2:**
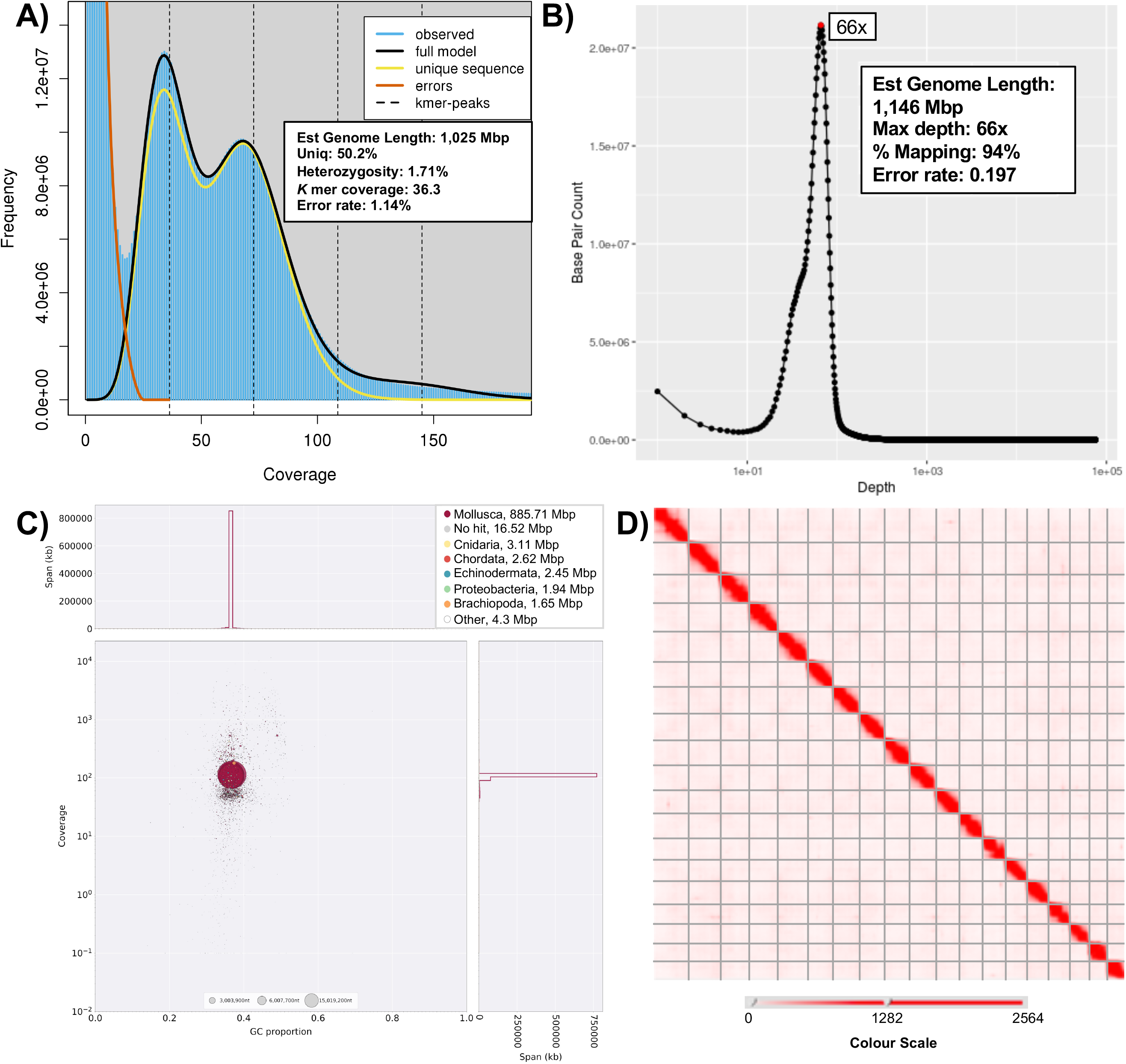
A) Genomescope2 [66] plot of the 21 mer *k-*mer content within the *Pecten maximus* genome. Models fitted and resulting estimates of genome size and read data as shown on figure. B) Basepair count by depth in PacBio data, determined using PBreads/Minimap2 C) Blobplot [42] of content of the *P. maximus* genome. Note little-to-no contamination of the assembly can be observed, with the small amount of sequence annotated as non-metazoan mirroring the metazoan content in GC content and average coverage. D) Hi-C contact map based on assembly created using 3D-DNA and Juicebox Assembly Tools (see http://bit.ly/2QaYqvk for an interactive version of this panel).

#### Assembly Assessment

The total size of our assembly, 918 Mbp, falls short of previous estimates of the genome size of *P. maximus*, with flow cytometry estimating a genomic c value of 1.42 [39]. Assessments of genome size based on *k-*mer counting using Genomescope (10,000 cov cutoff) [40] suggest that the complete genome size is approximately 1.025 Gbp (Fig. 2A). Estimates using PacBio reads and Minimap2 [41], showing basepair count at each depth, put the genome size at 1,146 Mbp, which is more in line with flow cytometry results. The reason for this discrepancy is likely to be caused by heterochromatic regions inaccessible to current sequencing technologies.

The expected genome size of *P. maximus* is slightly larger than many other sequenced bivalve species, and our assembly size (in bp) is in line with that of other sequenced scallop species (Table 3). It is, however, half the size of the genomes of the sequenced mussels *Gigantidas platifrons* and *Modiolus philippinarum.* Scallops therefore have intermediate genome sizes on average when compared to other molluscs, larger than oysters such as *Crassostrea gigas* and gastropods such as *Lottia gigantea*, but smaller than mussels and cephalopods.

To confirm the efficacy of the contamination screen performed during the assembly process, we verified the absence of parasitic or pathogenic sources by creating a Blobplot (Fig. 2C) using Blobtools [42]. We observed very few scaffolds (1.94 Mb, or around 0.21 % of our assembly) with blast similarity to Proteobacteria, but with coverage values and GC content exactly mirroring the rest of the assembly. In the majority of these cases, the assignment to Proteobacteria will be due to a chance blast match with high similarity over a small region of the contig length, rather than actual bacterial origin. The vast majority of the assembly (885.71 Mb) was assigned to the clade Mollusca, as expected (Fig. 2C).

To assay assembly quality and completeness, we mapped our raw reads to the genome. Of the 10x Genomics paired-end reads, 94% (814,387,200 of 866,234,784 reads) mapped concordantly. Of our PacBio reads, 94% (71.13 x10^9^ of 75.7 x10^9^ bases) also mapped (Fig 2B), indicating a well-assembled dataset, and one with little missing data.

The observed heterozygosity (1.71%, Fig. 2A) in the *P. maximus* assembly is a common phenomenon in broadcast spawning marine invertebrates [43]. It should be noted that we used Purge Haplotigs on our final assembly when using this resource for studies focusing on genetic diversity. Levels of heterozygosity in *P. maximus* were higher than those found in the Sydney Rock Oyster *Saccostrea* (0.51%), or the Pacific oyster *Crassostrea gigas* (0.73%). Both of these oyster samples were derived from selective breeding programmes, which would reduce heterozygosity compared to wild populations [20].

Repeat elements have been noted as playing an important role in genome evolution in molluscs, and in bivalves in particular [e.g. 44]. We used RepeatModeler and RepeatMasker [45] to identify and mask regions of the genome containing previously identified or novel repetitive sequences [Table 4]. With the caveat that not all repetitive elements have been classified, it seems that LTRs are less common in *P. maximus* compared to other species (0.52%, c.f. 1.35% in *S. glomerata* and 2.5% in *C. gigas*), but that SINES are more common (2.19%, c.f. 0.09% in *S. glomerata* and 0.6% in *C. gigas*). A total of 27.0% of the genome was classified as repetitive elements, with 16.7% of the genome made up of elements not present in preconfigured RepeatMasker libraries (but likely shared with other bivalve species). While the genome of *P. maximus* is large by scallop standards, its size is not due to large amounts of repetitive elements, as 27.0% is low compared to many other genome resources. For example, *Crassostrea gigas* has a repeat content of 36% [15] and *Saccostrea glomerata* 45.0% [20].

**Table 4:** Repeat content of the *P. maximus* genome based on RepeatModeler and RepeatMasker analysis.

| **Element** | **Count** | **Length Occupied**<br>**(bp)** | **% of Genome** |
| --- | --- | --- | --- |
| SINES | 125,121 | 20,067,275 | 2.19 |
| MIRs | 21,406 | 3,059,644 | 0.33 |
| LINES | 86,373 | 26,983,591 | 2.94 |
| LINE1 | 803 | 463,519 | 0.05 |
| LINE2 | 4,883 | 2,601,659 | 0.28 |
| L3/CR1 | 4,374 | 1,588,697 | 0.17 |
| LTR elements | 9,334 | 4,731,793 | 0.52 |
| DNA elements | 121,409 | 31,845,557 | 3.47 |
| hAT-Charlie | 1,312 | 394,533 | 0.04 |
| TcMar-Tigger | 4,548 | 1,478,364 | 0.16 |
| Unclassified | 612,341 | 153,700,734 | 16.74 |
| Total interspersed repeats |  | 237,328,950 | 25.84 |
| Small RNA | 4,096 | 563,615 | 0.06 |
| Simple repeats | 174,931 | 9,099,659 | 0.99 |
| Low complexity | 2,5658 | 1,411,700 | 0.15 |
| Total length:<br>(of 918.3 Mbp) |  | 247,513,725 | 27.0 |

#### Gene Prediction and Annotation

Gene sequences were predicted using Augustus annotation software [46], with one novel (K. James, available for download from Figshare link, see data sources section) and several previously published *P. maximus* RNAseq datasets [47, 48] used for training. The non-masked genome was used as the basis for gene prediction, to avoid artefacts, missed exons or missing gene portions caused by gene overlap with masked areas of the genome. UTR prediction, and gene prediction on both strands was set “true”, and two rounds of training (without and with UTR) took place.

This annotation resulted in an initial set of 215,598 putative genes (with 32,824 genes having two or more alternative isoforms, resulting in 249,081 discrete transcript models. We filtered the initial gene set by comparing our gene models to seven previously published bivalve resources using Orthofinder2, and retained genes with orthologues shared with other species (57,574 genes, further details below). To ensure we did not discard transcribed genes absent from other bivalves but present in our resource, we also retained those genes with a good hit in the nr database (23,541 genes, diamond blastp, --more-sensitive --max-target-seqs 1 --outfmt 6 qseqid sallseqid stitle pident evalue --evalue 1e-9), a total of 81,115 genes. However, we then removed from this combined total any genes which had a match within our identified repeat elements (13,374 genes, tblastn, -evalue 1.0e-29 -max_target_seqs 1 -outfmt ’6 qseqid staxids evalue’). This evalue cutoff was chosen after initial trials to include genes which mapped to *pol*, *env, tc3 transposase, Gag-Pol* and *reverse transcriptase* genes in automated blast. This resulted in a final, 67,741 gene, curated set, of which 16,693 genes possess one or more alternative transcripts. Full, curated and annotated gene sets in a variety of formats can be found at http://dx.doi.org/10.6084/m9.figshare.10311068. This number, while still high in comparison to the number of genes found in many metazoan species, is comparable to the number of unigenes (72,187) in the *Argopecten irradians* resource [49].

We assayed the completeness of our gene set using the BUSCOv2 (Benchmarking Universal Single Copy Orthologs, Simão et al 2015), using metazoan gene sets. Of the 978-gene Metazoa dataset, 924 (94.5%) complete BUSCOs (of which 32 (3.3%) were duplicated), 10 incomplete (1%) BUSCOs and 44 (4.5%) missing BUSCOs were recorded in genome mode, equating to a recovery of 95.5% of the entire BUSCO set. This is comparable to previously published bivalve resources, as can be seen in Table 3.

We have performed annotation of gene complements using a number of automated methods. BLAST annotation was performed using DIAMOND (--more-sensitive --max-target-seqs 1 --outfmt 6 qseqid sallseqid stitle pident evalue --evalue 1e-3 --threads 4), with 88,824 of our unfiltered gene models recovering a hit, although this figure includes hits to repetitive elements removed in our curated dataset (Supplementary File 2, Figshare). Of the 67,741 high confidence genes, 59,772 possess a hit in the nr database (88.2%), indicating a highly annotatable dataset. We also used the KEGG-KAAS automatic annotation server, using peptide sequence and the BBH method. The standard eukaryotic species set, complemented with *Lottia gigantea*, *Pomacea canaliculata*, *Crassostrea gigas, Mizuhopecten yessoensis* and *Octopus bimaculoides* was used for annotation, with 14,495 of our gene models mapping to KEGG pathways (Figshare, Supplementary File 3).

#### Gene complement and expansion

We investigated the gene complement of *P. maximus* to understand the nature of the events that resulted in it and other scallops possessing a large number of annotated genes compared to related mollusc species. This analysis was performed predominantly using Orthofinder2 (-t 8 -a 8 -M msa -T fasttree settings, and using only the longest transcript per gene for *P. maximus*) and shown in Figure 3A. These statistics reveal that *P. maximus* exhibits little gene loss compared to other related species. The percentage of orthogroups containing *P. maximus* genes is very high (83.4%) compared to every other species examined. *P. maximus* has therefore lost fewer genes from the ancestrally shared cassette than any of the other species listed. *Pecten maximus* also possesses 518 species-specific orthogroups – comparatively more than any other species listed. These genes are likely to be true novelties, as they are not found in any of the eight other species of bivalve examined here.

**Figure 3:**
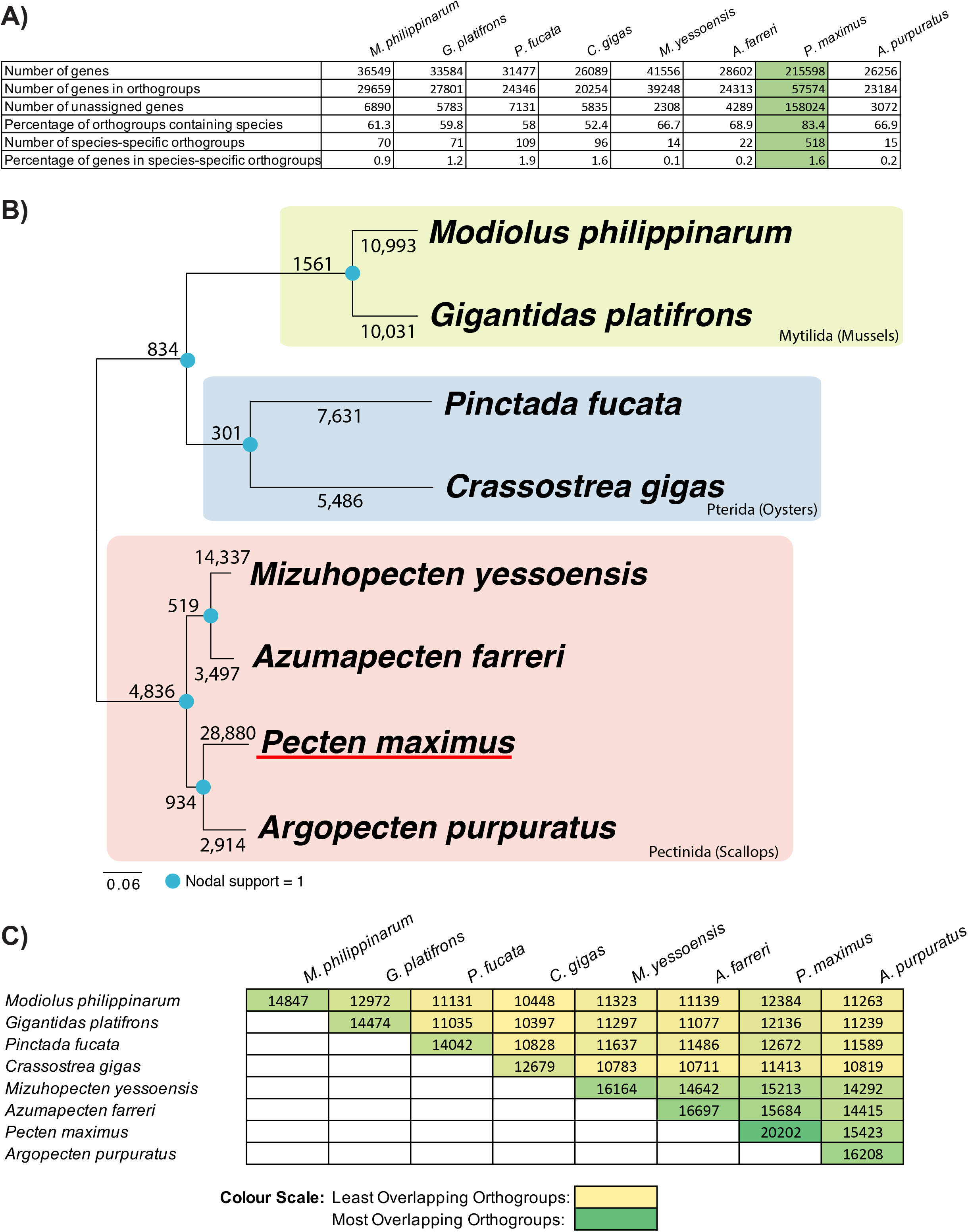
A) Orthofinder 2 [67] ortholog analysis of eight sequenced marine bivalve species. *Pecten maximus* results shown in green. B) Phylogeny of bivalves using available marine bivalve genomes (generated from ortholog groups by STAG and displayed in Figtree), with root placed at midpoint. Blue dots indicate nodal support (=1 at every node). Numbers on internal nodes represent ancestrally shared duplications at the point of diversification. Numbers on leaf nodes indicate duplication events occurring solely in that taxon. C) Matrix showing numbers of overlapping orthogroups shared by the species examined. A colour scale has been applied to aid in identifying the most- and least-overlapping data sources.

Using these results, we are also able to understand the prevalence of gene duplication across the phylogeny of bivalves. Figure 3B shows gene duplication events inferred from the orthogroup analysis as mapped onto the phylogeny of the eight bivalve species examined here. We conclude that gene duplication events are common in extant species of bivalve, and some gene duplicates are shared by leaf nodes as a result of events in the stem lineage. However, duplications in *P. maximus* are particularly prevalent. With 28,880 unique duplications, *P. maximus* has more than double the number of duplicates than any species, with *Mizuhopecten yessoensis* the next closest example. However, it should be noted that not all gene annotations were performed in an identical fashion, and particularly if genes have been missed in other species, for example through sparse RNAseq for gene prediction, this will negatively influence their counts in these results.

Of the genes that are shared with other lineages, *P. maximus* has a highly complete complement (Fig. 3C). No other species examined here possesses as many shared orthogroups in total, or shares as many with other species. In pairwise comparisons, only the mussels *Modiolus philippinarum* and *Gigantidas platifrons* show similar numbers of shared orthogroups with each other, but not with other species. This is consistent with the previous finding that the scallop *Mizuhopecten yessoensis* is closer in gene complement to the oysters *Crassostrea gigas* and *Pinctada fucata* than the oysters are to one another [18], a fact reflected in early divergence of these two distantly related oyster species [50]. Scallops in general therefore have a better-conserved gene cassette compared to the ancestral genotype than exhibited in oysters.

We conclude *P. maximus* has a well-conserved gene set, that has been added to substantially by gene duplication. Its large gene complement is therefore explained by a strong pattern of gene gain, coupled to very little gene loss.

#### Hox genes

The prevalence of gene duplication within *P. maximus* led us to consider whether a whole genome duplication (WGD) event had occurred in this lineage. As a test for this, we used the well-conserved Hox and Parahox gene clusters, which are normally preserved as intact complexes and duplicated in the presence of additional WGD events [e.g. 51, 52].

*Pecten maximus* possesses a single Hox cluster spanning 1.72 Mbp (from 28,829,013 bp– 30,558,725 bp) on scaffold HiC_scaffold_2_arrow_ctg1 (Fig. 4A). It also features a single Parahox cluster on scaffold HiC_scaffold_5_arrow_ctg1. The complex, like that of *Mizuhopecten yessoensis* [18], is stereotypical. This evidence suggests that no WGD has taken place.

**Figure 4:**
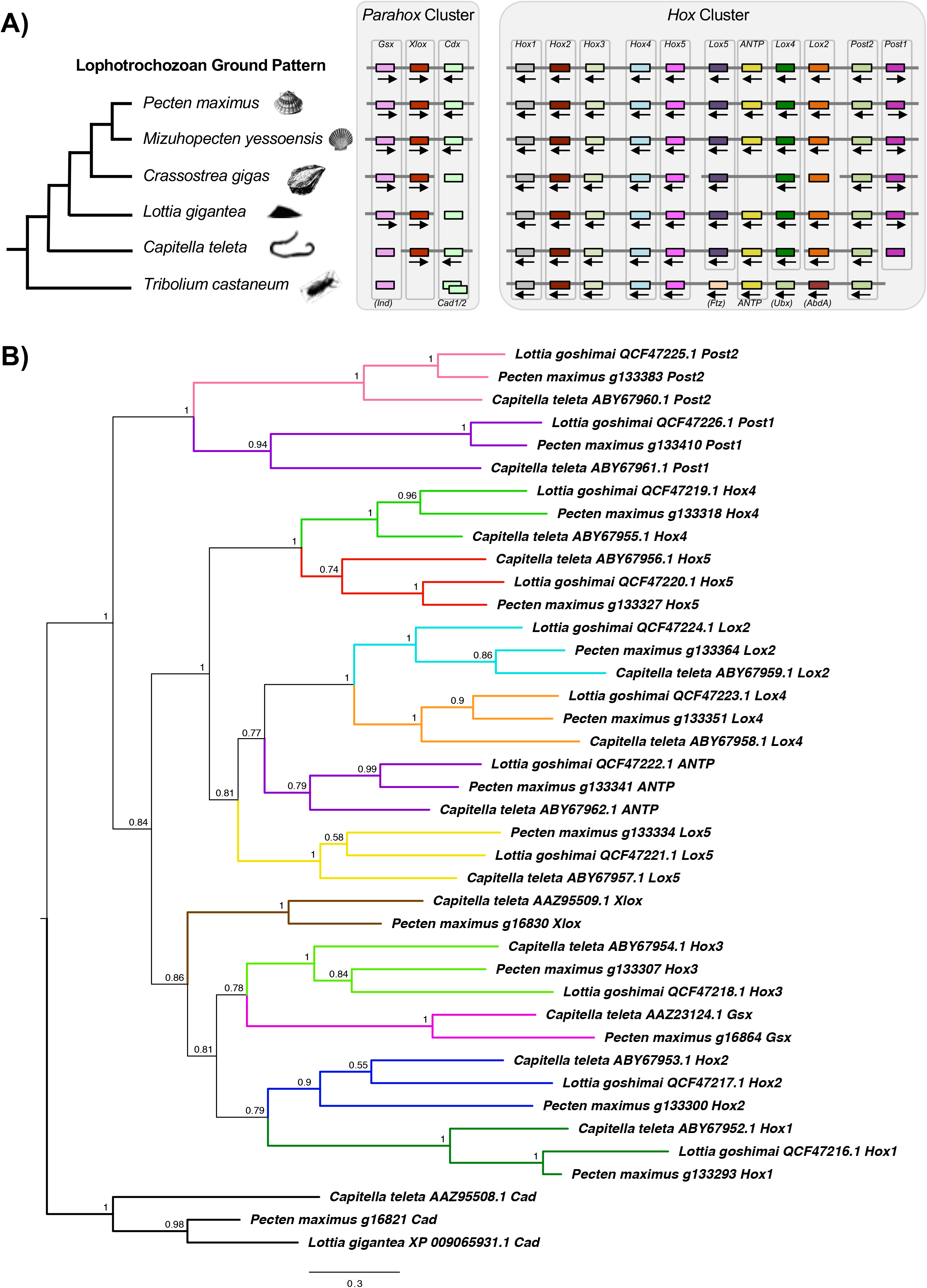
A) Diagrammatic representation of *Hox* and *Parahox* cluster chromosomal organisation showing a shared pattern among selected Lophotrochozoan taxa (scallops *Pecten maximus* and *Mizuhopecten yessoensis*, Pacific oyster *Crassostrea gigas*, owl limpet *Lottia gigantea* and annelid, *Capitella teleta*) along with an outgroup (red flour beetle; *Tribolium castaneum)*. Grey bar linking genes represents regions of synteny. Silhouette sources noted in Acknowledgements. Arrows show direction of transcription where known. B) Phylogeny of *P. maximus* Hox and Parahox genes alongside those of known homology from previous work [68, 69] inferred using MrBayes [70] under the Jones model (1,000,000 generations, with 25% discarded as ‘burn-in’). Numbers at base of nodes are posterior probabilities, shown to 2 significant figures. Branches are coloured by gene.

#### Immunity to neurotoxins

Bivalves are known to accumulate a number of toxins derived from phytoplankton, and human ingestion of contaminated bivalves can result in five known syndromes: Amnesiac Shellfish Poisoning (ASP) caused by domoic acid (DA), Paralytic Shellfish Poisoning (PSP) from saxitoxins (STX), Diarrhetic Shellfish Poisoning from okadaic acid and analogues, Neurotoxic Shellfish Poisoning caused by brevetoxin and analogues, and Azaspiracid Shellfish Poisoning from azaspiracid [12]. Adult *P. maximus* are relatively immune to STX and DA and as such, may be vectors for the syndromes PSP and ASP, which are of the greatest concern to human health [53, 54].

STX and brevetoxin are neurotoxins that bind to the voltage-gated sodium channel, blocking the passage of nerve impulses [55]. Previous studies have shown that genetic mutations within the sodium channel gene, *Neuron Navigator 1* (*Nav1*) confer immunity in taxa that accumulate saxitoxin (e.g. the soft-shell clam *Mya arenaria* [56]; scallop *Azumapecten farreri* [17]; copepods, *Calanus finmarchicus* and *Acartia hudsonica* [57]) or other similar acting neurotoxins like tetrodotoxin (TTX) (e.g. pufferfish, *Tetraodon nigroviridis* and *Takifugu rubripes*; salamanders [58–61]; and the venomous blue-ringed octopus [62]).

The *P. maximus Nav1* gene possesses the expected canonical domain structure observed in other taxa. Furthermore, it possesses the characteristic thymine residue in Domain 3 (Fig. 5, position 1425 in reference to rat sodium channel IIA), also described in the other two scallop species sequenced so far, which has been shown to confer resistance to these toxins in pufferfish, copepods and the venomous blue-ringed octopus [57–59]. It does not, however, have the E945D mutation seen in the softshell clam *Mya arenaria* and some pufferfish, which experimental evidence suggests also confers resistance [56], nor the D1663H or G1664S mutations in the blue-ringed octopus [62]. Instead, it has one novel and two ancestrally shared changes (shared with scallops and other bivalves) that may be of interest in studying alternative means of resistance in this molecule.

**Figure 5:**
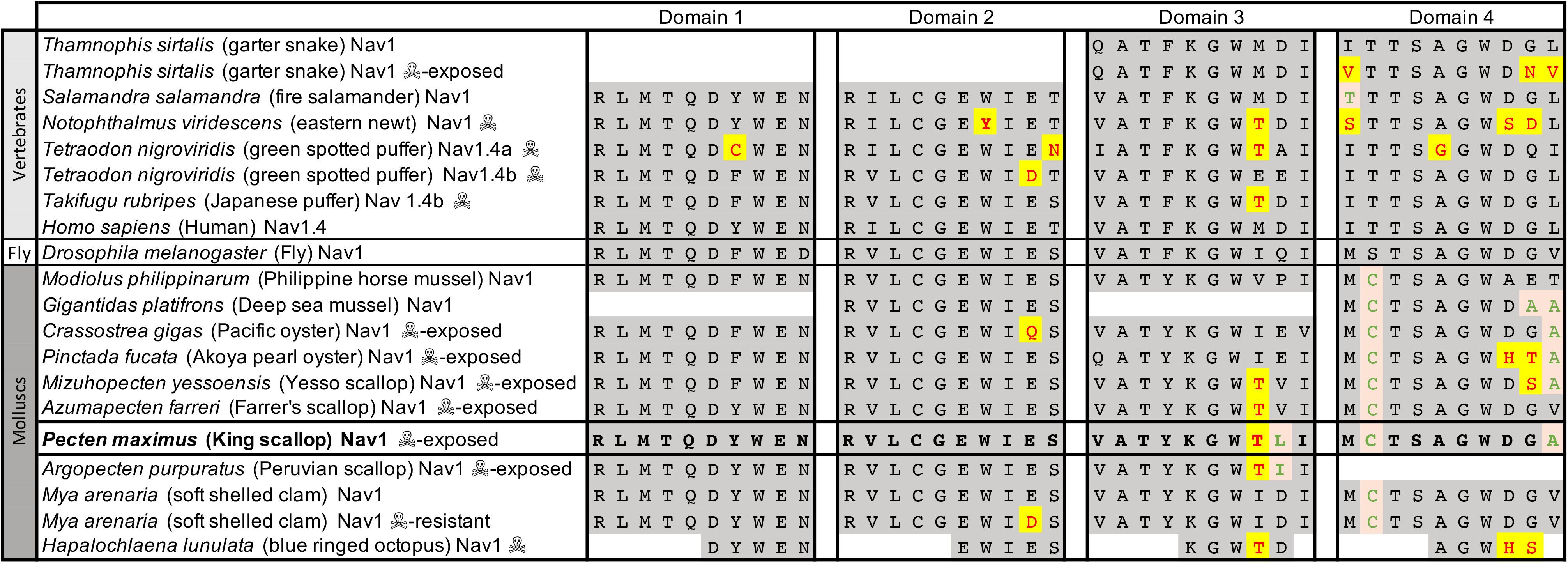
Domain alignments of the sodium channel *Nav1* showing residues (text in red, highlighted in yellow) implicated in resistance to the neurotoxins tetrodotoxin (TTX) and saxitoxins (STX). Species of vertebrate and mollusc known to be resistant to TTX or STX [58–61] are shown alongside species and sub-populations with no resistance to these toxins. Species (and sub-populations) that produce or accumulate these toxins with little or no ill effect are marked with a skull-and-crossbones. *Pecten maximus* (bold text) shares a thymine residue in domain 3 known to confer neurotoxin resistance in several other species. It also has a number of residues (shown in green text with amber background) in domain 3 and 4, which are either unique to *P. maximus* or shared with other resistant shellfish, but not seen in other species. These residues are good candidates for testing for a functional role in resistance in the future.

Unlike STX and TTX, DA does not directly target sodium channels, instead it mimics glutamate and binds preferentially to glutamate receptors including N-methyl-D-aspartate (NDMA), kainate and α-amino-3-hydroxy-5-methyl-4-isoxazolepropionic acid (AMPA) receptors leading to elevated levels of intracellular calcium and potentially, calcium toxicity [9, 63]. A recent study, however, has shown that extracellular sodium concentration plays a crucial role in excitotoxicity of DA [64], suggesting that mutations we observe at *Nav1* may also confer a degree of immunity to DA in *P. maximus*. This has ramifications for the study of neurotoxin resilience and prevalence in the increasingly important commercially fished populations of *P. maximus*.

#### Conclusions

The genome of *Pecten maximus* presented here is a well assembled and annotated resource that will be of utility to a wide range of investigations in scallop, bivalve and molluscan biology. It is, to date, the best scaffolded genome available for bivalves, despite the heterozygosity seen in this clade. Given this assembly is based on state-of-the-art long-range data and has undergone structural verification, this resource will be particularly key for comparative analysis of structural variation and long range synteny. The curated gene set of this species exhibits little loss compared to other sequenced bivalve species, and possesses numerous duplicated genes which have contributed to the largest gene set observed to date in molluscs. The genes are well-annotated, with 88.2% of our high confidence gene set mapped to a known gene. This genome has already yielded a range of insights into the biology of *P. maximus*, and will provide a basis for investigations into fields such as physiology, neurotoxicology, population genetics and shell formation for many years to come.

## Declarations

### List of abbreviations

AMPA: α-amino-3-hydroxy-5-methyl-4-isoxazolepropionic acid receptors
ASP: Amnesiac Shellfish Poisoning
BLAST: Basic Local Alignment Search Tool
BUSCO: benchmarking universal single copy orthologs
DA: domoic acid
LINES: Long Interspersed Nuclear Elements
LTRs: Long Terminal Repeats
MIRs: Mammalian Wide Interspersed Repeats
NDMA: N-methyl-D-aspartate receptors
PSTs: paralytic shellfish toxins
STX: saxitoxins
SINES: Short Interspersed Nuclear Elements
TTX: tetrodotoxin
UTR: Untranslated Region
WGD: whole genome duplication

### Ethics approval and consent to participate

Not applicable

### Consent for publication

Not applicable

### Authors’ contributions

STW conceived of the study, provided the tissue samples and contributed to the text. NJK performed bioinformatic analyses, drafted the manuscript and prepared the figures. SAM assembled the draft genome. OD, ADO, DW and ELA generated and analysed the Hi-C data as part of the DNA Zoo effort. YR and KJ contributed to bioinformatic analyses, particularly RNAseq. KH lead the assembly curation, with JT performing contamination checks and removal, YS creating assembly analyses and SP performing manual assembly curation. EB, CC, JD KO, JS, MS and AW aided with DNA extraction, processing, sequencing and data delivery. DM and KH were responsible for project organisation. All authors approved the final version of the manuscript.

## Acknowledgements

The authors wish to thank the members of the Riesgo and Williams lab groups for helpful discussions in preparation of this resource. We thank Phylopic, and particularly B. Duygu Özpolat and Taro Maeda (http://creativecommons.org/licenses/by-nc-sa/3.0/) for images in Fig 4. Other images from the public domain include: *Pecten maximus* from Gosse*: Natural History: Mollusca* (1854). *Tribolium castaneum* from Comstock: *A manual for the study of insects* (1895). Oyster from Lear: Alphabet of Nonsense. Scallop from Popular Science Monthly, Vol 49, 1896.

## Data Statement

The *Pecten maximus* xPecMax1.1 assembly is available under the accession GCA_902652985.1. The data sets supporting the results of this article are available from FigShare, with accession number http://dx.doi.org/10.6084/m9.figshare.10311068 and on the DNA Zoo website at www.dnazoo.org/assemblies/Pecten_maximus.

## Additional Files

**Supplementary File 1:** Read quality assessment, FastQC/NanoComp. Zipped html files.

**Supplementary File 2:** BLAST annotations, *Pecten maximus* gene models. Zipped text files.

**Supplementary File 3:** KEGG-KAAS annotations, *Pecten maximus* gene models. Zipped text files.

