## Supplementary figures and images for "The Gene-Rich Genome of the Scallop *Pecten maximus*"

### NanoComp_lengths.pdf

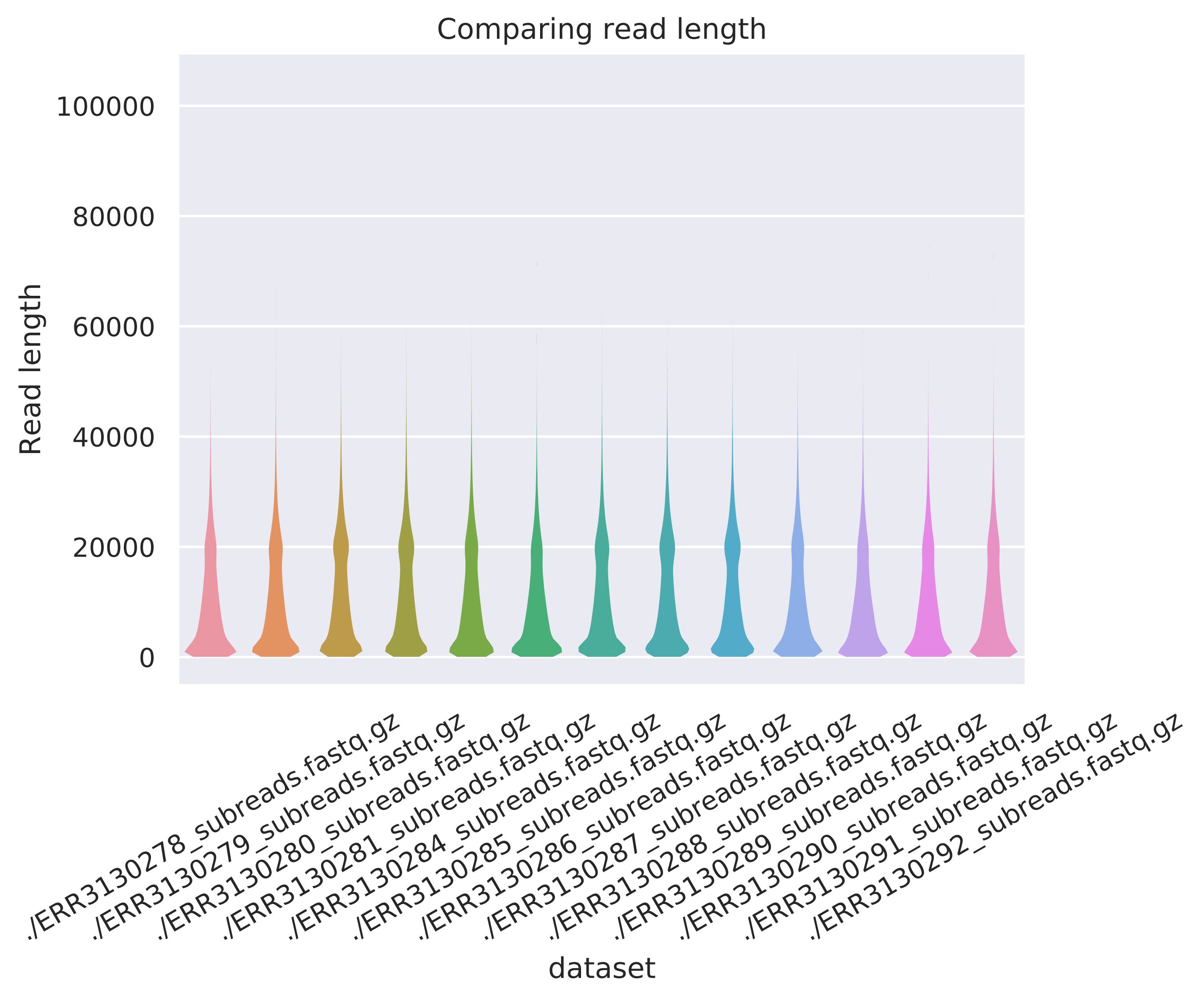

### NanoComp_log_length.pdf

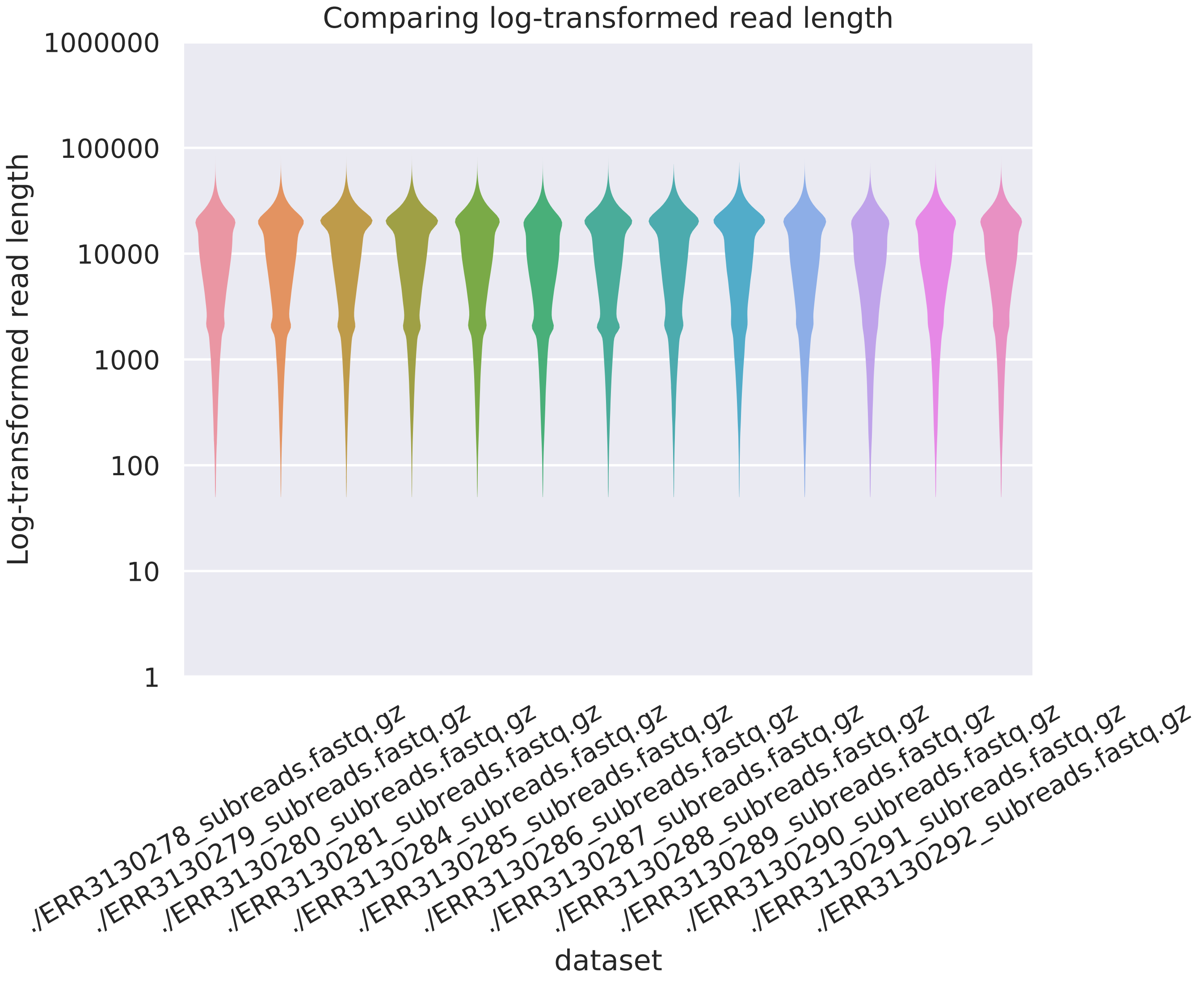

### NanoComp_N50.pdf

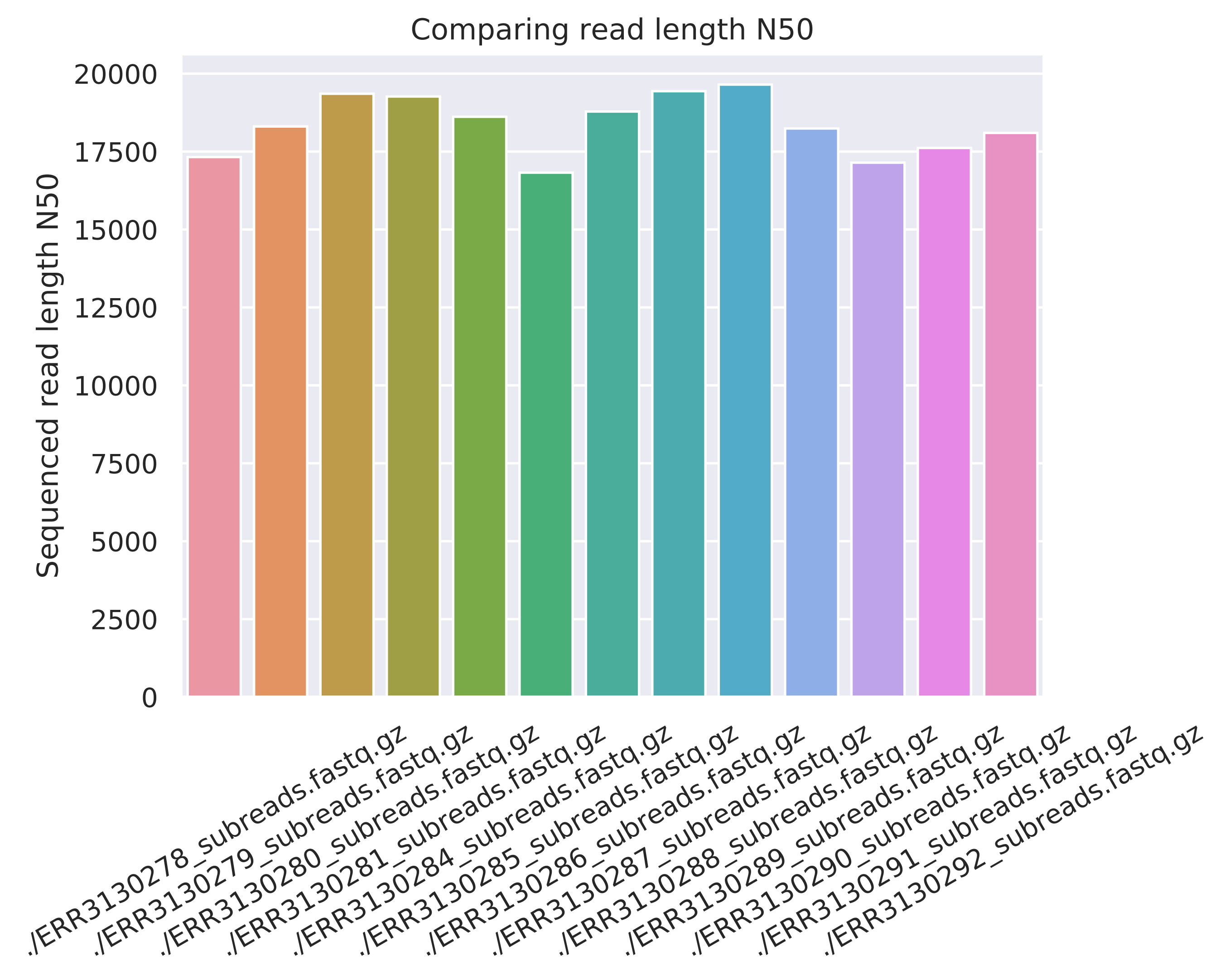

### NanoComp_number_of_reads.pdf

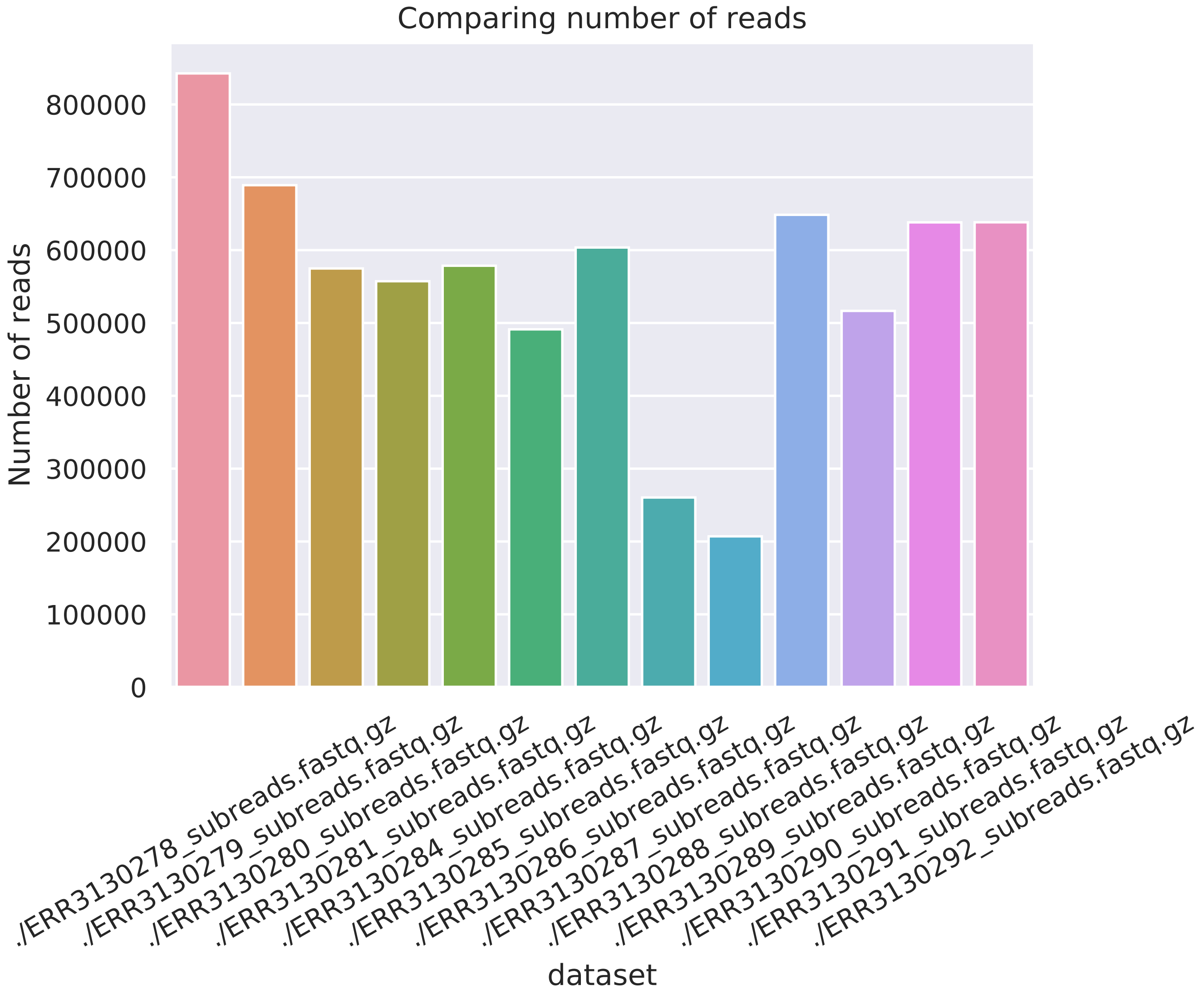

### NanoComp_total_throughput.pdf

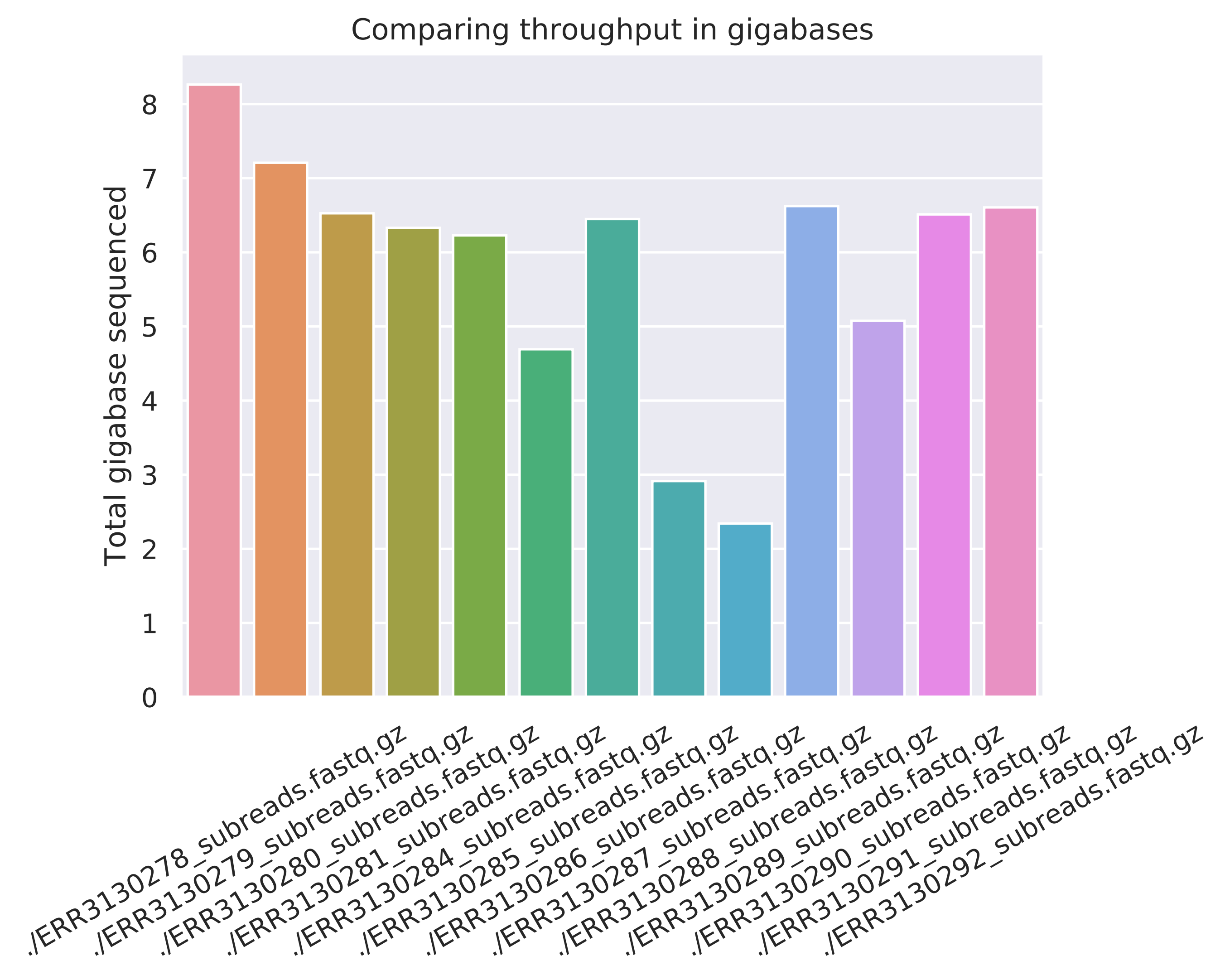
